# Quantification of cancer cell migration with an integrated experimental-computational pipeline

**DOI:** 10.1101/130526

**Authors:** Edwin F. Juarez, Carolina Garri, Ahmadreza Ghaffarizadeh, Paul Macklin, Kian Kani

## Abstract

We describe an integrated experimental-computational pipeline for quantifying cell migration *in vitro.* This pipeline is robust to image noise, open source, and user friendly. The experimental component uses the Oris cell migration assay (Platypus Technologies) to create migration regions. The computational component of the pipeline creates masks in Matlab (MathWorks) to cell-covered regions, uses a genetic algorithm to automatically select the migration region, and outputs a metric to quantify the migration of cells. In this work we demonstrate the utility of our pipeline by quantifying the effects of a drug (Taxol) and of the secreted Anterior Gradient 2 (sAGR2) protein in the migration of MDA-MB-231 cells (a breast cancer cell line). In particular, we show that blocking sAGR2 reduces migration of MDA-MB-231 cells.

## Introduction

### Objectives

We set out to design and implement a pipeline to quantify cell migration that is robust to image noise, open source, and user friendly.

### Background

In order to understand and treat cancer, we need to study and ultimately control metastasis [1]. A key aspect of metastasis is cell migration [1]. The anterior gradient protein 2 (AGR2) has been shown to promote cell migration [2]. High expression of AGR2 is correlated with aggressive forms of various adenocarcinomas including prostate [3] and breast cancer [4]. Therefore, AGR2 could be used as a biomarker and therapeutic target for cancer.

In this work, we describe an experimental and computational pipeline to quantify cell migration. We demonstrate this pipeline by quantifying migration of MDA-MB-231 cells, a breast cancer cell line known to migrate aggressively [5]. We show that blocking the secreted AGR2 (sAGR2) with an antibody that binds specifically to AGR2 (referred to as AGR2-Ab) in cell medium prevents the migration of the MDA-MB-231 cells. Our pipeline aids in the verification of well-established hypotheses and it can be used to test new hypotheses, thus, aiding in and accelerating the drug discovery process.

## Methods

### Cell migration assay

The Oris^TM^ migration assay use a physical barrier “stopper” to create a defined circular region that is intended prevent cell adhesion at the start of the assay. This central cell-free detection zone is in the center of each well of a 96-well plate. As the cells migrate to the cell-free zone over 24-48 hours, real-time assessment of migratory cells allows acquisition of richer data sets. Since there are no artificial membranes or inserts in the light path through which cells must pass, this assay is amenable to quantification with microscopy. We used the Oris cell migration assay from Platypus Technologies to create migration regions by inserting stoppers in each of the 96 wells on a plate. Shortly after inserting the stoppers, we seeded MDA-MB-231 cells and waited until they reached 80% confluent (approximately 24 hours). Next we removed the stoppers and allowed the cells to move into the migration region. 48 hours after removal of the stoppers, we imaged each well using a fluorescence microscope.

### Automatic selection of migration region

We first created a mask corresponding to the area covered by cells using standard deviation filtering and applying a series of morphological operations in Matlab R2016a (MathWorks) as shown in figure 1.

**Figure 1.**
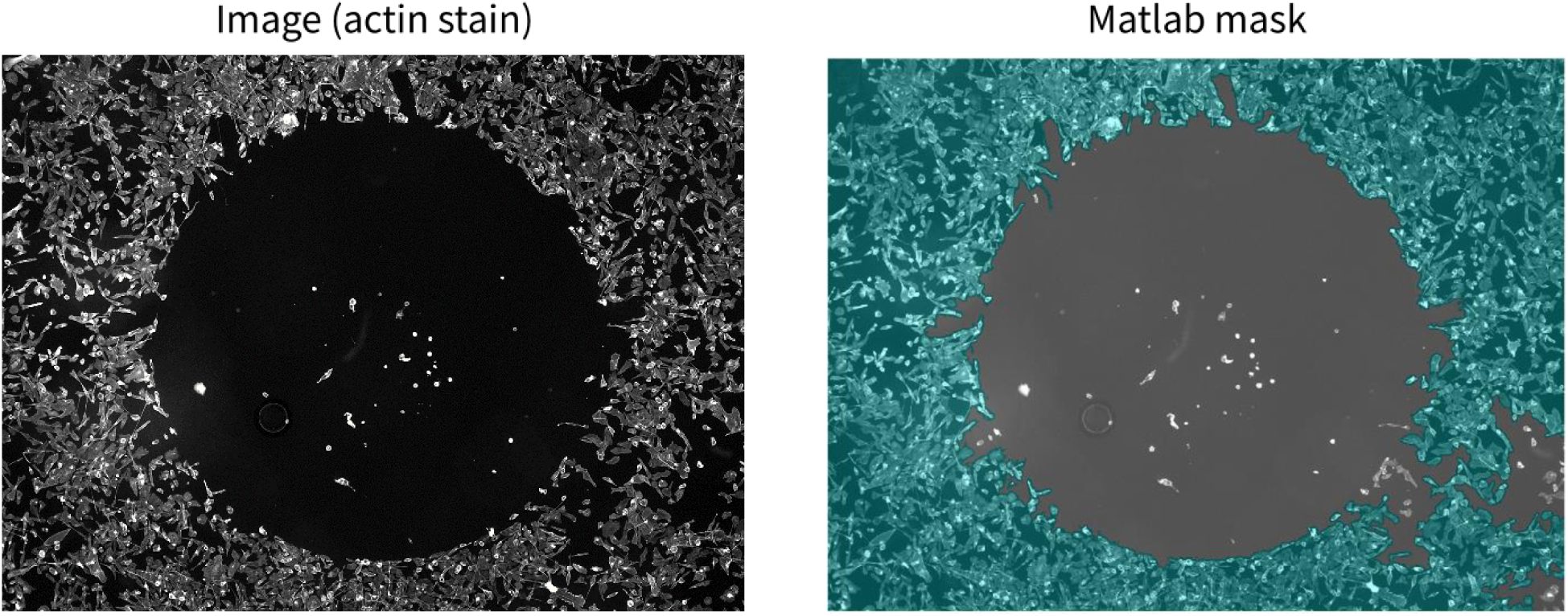
Sample images immediately after the stopper was removed. In order to identify the migration region, we took images of each well (left), then we select a mask that covers the area utilized by cells, highlighted in green (right).

Note that these images are black and white (green is used throughout to highlight software outputs), hence every pixel’s value belongs to the interval [0,1] where a completely black pixel has value 0 and a completely white pixel has value 1. Also note that a mask is a binary matrix that indicate which pixels belong to the mask (with value of 1, these pixels are referred to as “cell pixels”) and which pixels do not belong to the mask (with value 0).

We then used a genetic algorithm to determine the coordinates of the center and the radius of a circle according to equation (1). This optimal circle determines the migration region.

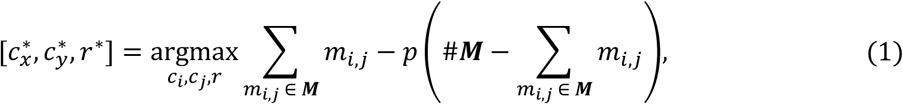

where 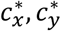, and *r*\* are the optimal parameters of the migration region, ***M*** is the Matlab mask we are evaluating (i.e., a circle with center at coordinates (*c_x_, c_y_*) and radius r), so 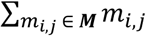 is the sum of all the pixel intensities (my) which belong to the mask ***M***, ***#M*** is the cardinality of ***M*** (i.e., the number, of pixels which belong to ***M***), and *p* is a penalty parameter. If *p* = 1 we have:

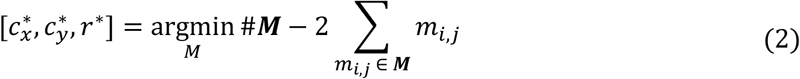

Hence, the maximization problem from equation (1) is equivalent to the minimization represented in equation (2) (when *p* = 1). From equation (2), we can interpret the optimization performed by the genetic algorithm as finding “the largest circle which contains the least number of cell pixels.” Figure 2 shows the optimal circular region selected by the genetic algorithm when the input is the image from figure 1.

**Figure 2.**
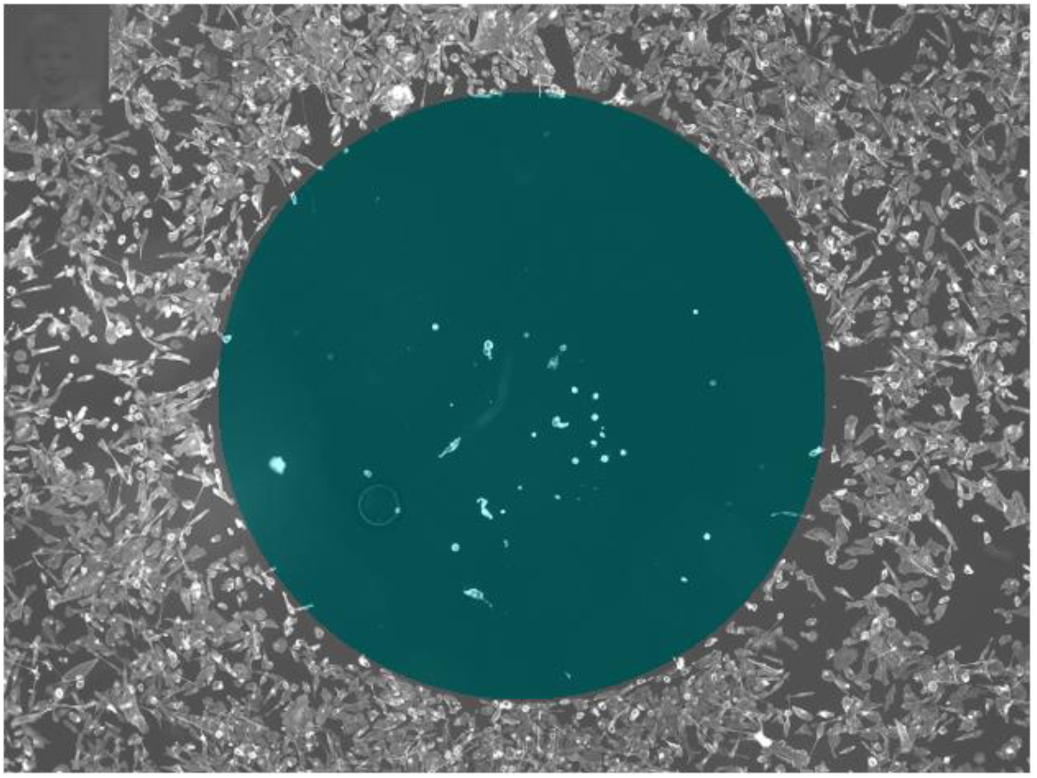
Automatically-selected migration region. The genetic algorithm selects the largest circle which contains the least number of masked pixels from the cell area mask. This optimal circle is the migration region.

### Percent of migration region covered by cells

We defined a metric to quantify the migration of MDA-MB-231 cells. This metric is *Q,* the percentage of migration pixels inside the migration region. We define a migration pixel as any pixel whose intensity value is greater than or equal to a threshold *T.* We chose *T* equal to 1.25 times the median pixel intensity of the migration region immediately after the stopper was removed (i.e., the green region in figure 2). This is:

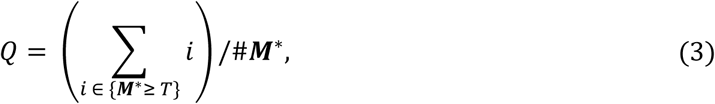

Where ***M\**** is the optimal circle defined by the three parameters (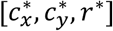 from equations (1) and (2)) and the set {***M\**** *≥ T*} includes all of the pixels inside ***M\**** with intensities higher than *T.*

## Results and Discussion

To test the hypothesis that MDA-MB-231 cells’ migration is reduced in the absence of AGR2, we designed an experiment with 5 experimental conditions: a positive control (untreated cells), a negative control (wells where the stopper was not removed), cells treated with lOnM of Taxol (a non-cytotoxic dose level which prevents cell migration but not cell death [6]), cells treated with a 1:50 dilution of AGR2-ab to inactivate sAGR2, and with a 1:50 dilution of IgG, a control antibody (Ctrl-Ab) which is does not affect cell migration. Representative images from these conditions (i.e., replicate 3) are shown in figure 3 (top).

**Figure 3.**
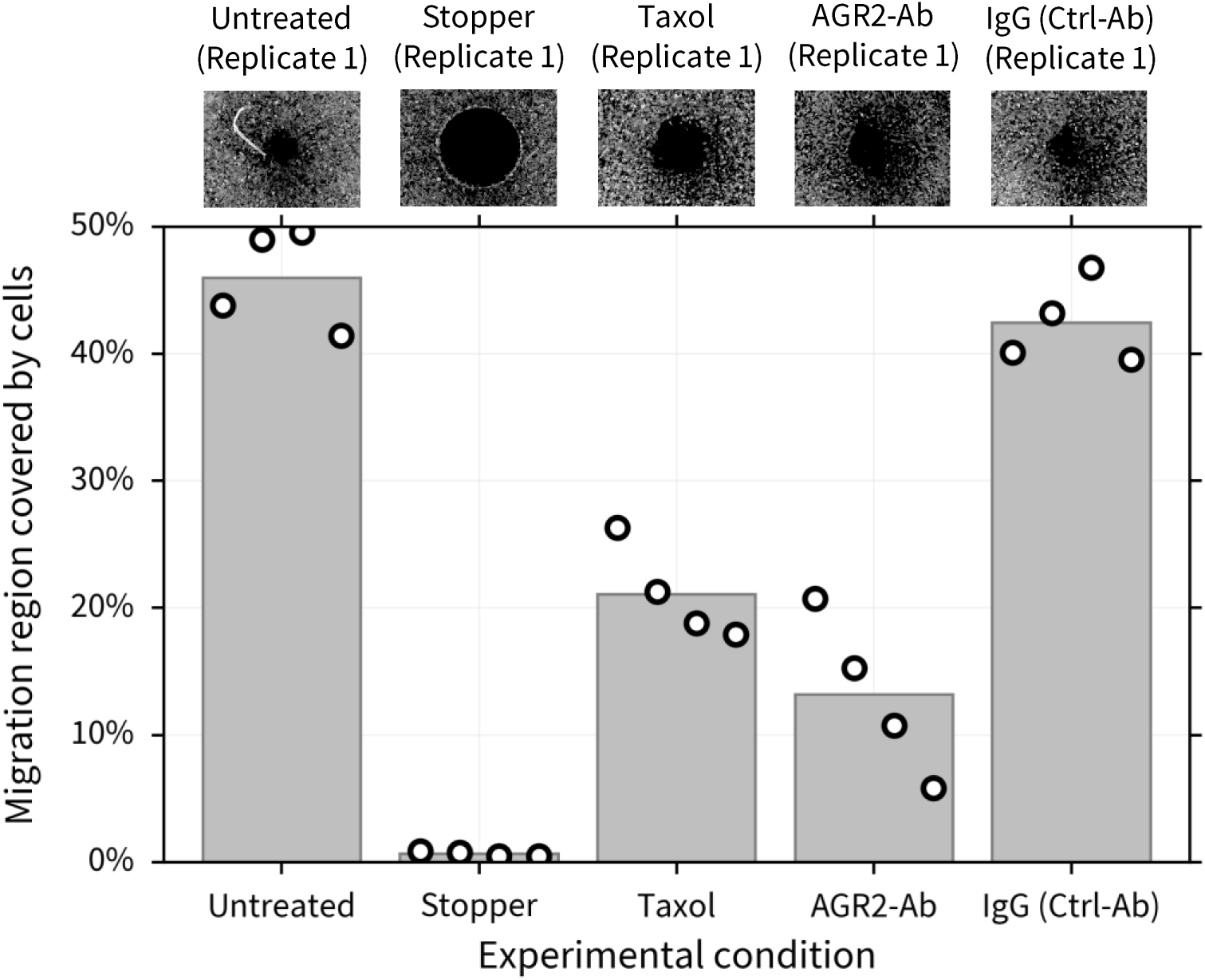
Quantifying cell migration. Representative images (replicate 3) of each condition are shown (top). lOnM of Taxol and thel0|ig/mL of the H10 peptide show similar levels of migration inhibition compared to the positive and negative controls. Our metric (bottom) allows us to quantify the qualitative results (top).

For the untreated case and the control peptide we observe high levels of migration, with a 45.96±1.99 (mean ± standard error of the mean) percent of the migration region covered in the untreated case and 42.44±1.66 percent of the migration region covered in the control antibody case. We fail to reject the null hypothesis that these two are the sample means from the same distribution (p value of 0.084). In the Taxol case, 21.07±1.88 percent of the migration region is covered. We reject the null hypothesis that the mean of the Taxol population and the mean of the untreated case are sample means from the same distribution (p value of 8.16e-5). Similarly, for the AGR2-Ab case, 13.15±3.18 percent of the migration region is covered. We reject the null hypothesis that the mean of the AGR2-Ab population and the mean of the untreated case are sample means from the same distribution (p value of 2.19e-5). Not only we confirm the hypothesis that MDA-MB-231 cells’ migration is reduced in the absence of AGR2, but our method allows for reproducible quantification of qualitative observations.

Furthermore, the algorithms used to compute *Q* require a single input from the user (a string with the names of the control experiments) and it can run in a desktop machine with Matlab (R2015a and above) installed.

## Conclusions

We have designed and implemented a pipeline for quantifying cell migration *in vitro.* This pipeline is robust to image noise, open source, and user friendly. In particular, we show that H10 aids in the reduction of migration of MDA-MB-231 cells by blocking sAGR2.

## Acknowledgements

We thank the USC Center for Applied Molecular Medicine for generous resources, the National Institutes of Health (Physical Sciences Oncology Center grant 5U54CA143907 for Multi-scale Complex Systems Transdisciplinary Analysis of Response to Therapy (MCSTART), and 1R01CA180149), the Breast Cancer Research Foundation, the USC James H. Zumberge Research and Innovation Fund, and USC Provost’s PhD fellowship for their generous financial support.

## Contributions

Conceptualization of the main idea presented in the paper: EFJ, CG, AG, and KK. Model development: EFJ, CG, and AG. Software & Computation: EFJ and AG. Biological Experiments: CG and KK. Analysis of the results: EFJ, CG, and AG. Writing - Original Draft: EFJ. Writing - Review & Editing: EFJ, CG, AG, PM, and KK. Supervision: PM and KK.

